# Neuronal ramping and theta power during freely-moving and head-fixed mouse interval timing

**DOI:** 10.64898/2026.05.27.728250

**Authors:** Matthew. A. Weber, Jacob Rysted, Kalpana Gupta, Alexandra S. Bova, Peter J. Bosch, Young-cho Kim, Nandakumar S. Narayanan, Georgina M. Aldridge

## Abstract

An important paradigm to study executive function is interval timing, which requires participants to estimate a temporal interval, often with a motor response. Interval timing translates from rodents to humans and can model neurodegenerative and neuropsychiatric disease. Interval timing also has parallel, time-dependent neurophysiology in rodents and humans, including time-dependent linear changes in neuronal firing rates over a temporal interval (ramping activity) and low-frequency ∼4 Hz “theta” activity evoked by trial start and response. Despite these translational features, an important confound of interval timing is movement and motor preparation, as participants must report their estimates by planning a movement. To address this confound, we first trained a group of 7 mice in a freely-moving interval timing task in which mice had to move across an operant chamber and nosepoke at the correct time to receive food reward. These mice were then trained in a head-fixed version of interval timing in which mice receive liquid reward for holding still for 3 seconds at the correct location, with reward access randomized to probe interval timing behavior. Despite vastly different motor sequences, we found prominent ramping activity in mouse prefrontal cortex ensembles and prefrontal cortical ∼4 Hz activity evoked both by trial start and preceding the timed decision. These data provide evidence that prefrontal neuronal ramping and theta activity is not linked to a specific motor program but rather a feature of the temporal organization of behavior.

## INTRODUCTION

Executive functions are high-level cognitive processes that promote goal-directed behavior. (Diamond, 2013). These functions include working memory, attention, and timing as well as processes such as inhibiting impulsive behaviors and engaging in selective attention (Fuster, 2001). Executive dysfunction is common in many neurologic and psychiatric diseases, including Alzheimer’s disease, schizophrenia, and substance abuse, and often is the characteristic pattern of cognitive decline in Parkinson’s disease and Lewy Body Dementia (Albert, 1996; Andreasen, 1999; Dolan et al., 2008; Parker et al., 2013). A highly translational paradigm to investigate the neural bases of these problems is interval timing, which is a cognitive process that requires participants to track the passage of time over seconds-to-minutes (Merchant & De Lafuente, 2014). Humans with these diseases have deficits in interval timing (Albert, 1996; Baddeley et al., 2001; Caselli et al., 2009; Malapani et al., 1998; Parker et al., 2017; Penney et al., 2005; Singh et al., 2021; Stopford et al., 2012; Wittmann et al., 2007) that can be modeled in rodents (Balci, Ludvig, et al., 2008; Buhusi & Meck, 2005; Parker et al., 2017; Ward et al., 2011; Weber et al., 2023, 2025).

Work from our group and others have identified time-dependent linear changes in firing rates over an interval as a key pattern of activity in which the prefrontal cortex encodes temporal information (Emmons et al., 2017; Henke et al., 2021; Hinton & Meck, 2004; J. Kim et al., 2013; Y.-C. Kim et al., 2017; Parker et al., 2014; Weber et al., 2025). Importantly, prefrontal cortex ramping activity is not readily explained by movement velocity, response, or reward anticipation (Bova et al., 2026), suggesting that this pattern of activity is related to estimating the passage of time. However, most interval timing procedures in mice require movement to reflect temporal estimates and receive a reward, and it is unknown if movement is required to activate prefrontal cortex circuits and ramping activity. This led to the hypothesis that if time-dependent ramping activity is encoding temporal information, prefrontal cortex neuronal ensembles will display similar patterns of activity in an interval timing task without movement.

To test this hypothesis, we first recorded from mouse prefrontal cortex neuronal ensembles during a freely-moving operant-based interval timing ‘switch’ task used extensively in our lab (Bova et al., 2026; Bruce et al., 2021, 2025; Ding et al., 2026; Larson et al., 2022; Stutt et al., 2024; Weber et al., 2023, 2025) that requires movement and a motor response to receive a reward. We then trained and recorded from prefrontal cortex neuronal ensembles from the same mice on a novel head-fixed interval timing task that requires mice to estimate an interval of time by holding still to acquire a reward. We find evidence of similar patterns of ramping activity and evoked theta activity in both tasks, although nuances between tasks could reflect important behavioral differences and neuronal mechanisms guiding behavior.

## Methods

### Mice

All procedures were approved by the Institutional Animal Care and Use Committee (IACUC) at the University of Iowa (Protocol #3052039). We included a total of 7 mice (4 C57BL/6J (3 females, 1 male) and 3 DAT^IRES^-Cre (1 female, 2 males)). During freely-moving interval timing “switch” task training, mice were food restricted to 85% of their starting bodyweight, had access to *ad libitum* water, and were housed individually post-surgery, according to methods described in detail previously (Bova et al., 2026; Bruce et al., 2025; Weber et al., 2025). Mice were implanted with recording electrodes and allowed to recover (1 week), trained on freely-moving interval timing procedures (4 weeks), and neurophysiological data was collected in freely-moving interval timing sessions (2 weeks). Mice were then placed back on *ad lib* access to standard rodent chow and rested for several months. At the beginning of head-fixed training, mice were then switched to mild water restriction (*ad lib* water restricted to a 2-hour daily interval), as sucrose-water reward is used for this task (see details). Mice were then trained in the head-fixed interval timing task (2 months) and neurophysiological data was collected (1 month), prior to histology.

### Stereotaxic surgery

Prefrontal electrodes were implanted according to procedures described in detail elsewhere (Bova et al., 2026; Weber et al., 2025). Briefly, mice were anesthetized using 4.0% isoflurane delivered at 400 mL/min and then maintained under 1.5%–3.0% isoflurane at 120 mL/min (SomnoSuite, Kent Scientific, Torrington, CT, USA) for the duration of the surgical procedures. Craniotomies were drilled over the prefrontal cortex (AP +1.8, ML ±0.3) counterbalanced across hemispheres (left vs. right). Multielectrode recording arrays (4×4, 1 mm²; MicroProbes, Gaithersburg, MD) were implanted at a depth of DV −1.8 mm. Additional craniotomies were drilled to implant skull screws for grounding electrodes and securing headcap assemblies. All craniotomies were sealed using cyanoacrylate adhesive (“SloZap,” Pacer Technologies, Rancho Cucamonga, CA), accelerated with “ZipKicker” (Pacer Technologies), and reinforced with methyl methacrylate (AM Systems, Port Angeles, WA). After surgery, mice were given one week to recover before resuming food restriction and interval timing training.

### Freely-moving interval timing task

Mice were trained in standard sound-attenuated operant chambers (MedAssociates, St. Albans, VT) in a freely-moving interval timing ‘switch’ task described in detail previously (Balci, Papachristos, et al., 2008; Bova et al., 2026; Bruce et al., 2021; Weber et al., 2025). Briefly, each session consisted of approximately 50% “short” trials with a 6-second interval and 50% “long” trials with an 18-second interval. Both trial types used identical auditory and visual cues and were self-initiated by the mouse via a nosepoke at the rear of the chamber. During each trial, mice were required to respond at a designated nosepoke port corresponding to the trial type — short or long — after the appropriate interval to receive a 20-mg sucrose pellet reward (BioServ, Flemington, NJ).

On short trials, mice responded at the short nosepoke port and were rewarded for their first response after 6 seconds; these trials were not analyzed. On long trials, a reward was delivered for the first response at the long nosepoke port following 18 seconds. Since the cues were identical for both trial types, the optimal strategy was for mice to begin by responding at the short nosepoke port and then switch to the long nosepoke port if no reward was received after approximately 6 seconds. This transition — termed the *switch response* — was defined as the moment the mouse left the short nosepoke port and served as a measure of the animal’s internal estimate of time, consistent with other interval timing paradigms. Only long trials in which mice executed an appropriate switch were included in the analysis. Extensive detail on task-specific movements, rewards, infrequent errors can be found in our prior work (Bova et al., 2026; Bruce et al., 2025; Ding et al., 2026). After reaching stable performance on the task, typically within four weeks, mice were acclimated to handling and the neurophysiological recording procedures described under “Neuronal Recordings”.

### Head-fixed adaptation and habituation

After the freely-moving interval timing task, mice were removed from food restriction and fed *ad lib.* Mice were placed under anesthesia briefly as described above to attach a head plate (Neurotar Model 13, ferromagnetic) to the existing implant, allowing compatibility with the head-fixed interval timing task. The Neurotar homecage is a free-floating arena that allows mice to freely move to different parts of the arena while fixed in place. Mice were handled and habituated to head-fixed procedures similar to our previous work (Keyes et al., 2021). A standardized habituation plan was followed to ensure all mice received similar introduction to the Neurotar head-fixed system. On the first day of habituation, these already well-acclimated mice were held near the Neurotar cage for 5-10 minutes, with breaks, observing for normal behaviors (grooming, sniffing, exploring) and absence of freezing behavior. When normal behaviors were observed, they were placed into the carbon fiber enclosure without fixation to explore for 5-10 minutes, first with the pressurized air turned off, then turned on. On the second day, the mouse was placed in the head-fixed apparatus. The mouse can then walk in all directions “within” the floating carbon fiber cage. Habituation to head-fixation occurred over two days, lasting 20 minutes to 1 hour per habituation session, depending on the behavioral response of each mouse. A fully habituated and trained mouse requires no restraint to be placed on the Neurotar magnetic system. Instead, mice are guided under the head plate fixation and once the magnets secure, the mouse can walk freely on the floating platform (while their head stays in one place). Following habitation, mice began training on the head-fixed interval timing task described under “Neuronal Recordings”.

### Head-fixed interval timing task

We developed the STOP (Select Target by Optimal Pause) task to train mice to use a timed-pause, rather than movement, to indicate a choice. While the STOP mechanism can be used for evaluating multiple behaviors, by design, the timed-pause is ideal for testing whether time-dependent prefrontal cortex neuronal activity is similar when no movement is required during the elapsed interval. The mouse pauses on one of two targets, initiating a continuous audible tone that lasts for as long as the mouse remains still. On the rewarded side, the mouse receives a reward at 3 seconds and the trial ends. If the mouse is on the unrewarded target, the tone continues past 3 seconds, acting as a probe trial. Ideal behavior is moving to the opposite target when the optimal pause length (3 seconds) does not produce a reward.

This task is distinct from the freely-moving interval timing task in several ways: 1) a mouse ‘response’ is indicated by lack of movement, or a ‘pause’, rather than a nosepoke response; 2) the trained interval is 3 seconds, rather than 6 seconds, and the interval elapses only while the mouse is still; 3) the optimal response (pause) is 3 seconds at both target zones; and 4) the rewarded target zone (rather than interval length) is pseudo-randomized between trials.

In the head-fixed interval timing STOP task, a pause is defined as velocity less than 50 cm/s from the Neurotar tracking software. Airflow to the Neurotar is adjusted depending on mouse weight, such that the speed threshold is only exceeded when the mouse makes a defined footfall or step. Behavioral shaping took approximately 4-8 weeks and included four stages.

Stage 1 (minimum 1 month): A pause of more than 0.5 seconds on the initiation zone turns off the chamber lights, delivers a small sucrose reward from a mounted lick spout, and starts the trial. During this phase, a second pause in either target zone triggers an audible tone and if the mouse is on the correct target, a reward is given. If the mouse continues to pause in the rewarded target during this stage, additional rewards are given at a rate of 1/second for 3 seconds total.

Stage 2 (minimum 1 month): Identical to Stage 1 except that no reward is given for trial initiation.

Stage 3 (minimum 1 month): Identical to Stage 2 except that the pause must be greater than 1 second to receive a reward.

Stage 4 (minimum 1 month): Identical to Stage 3 except that the mouse must pause for 3 seconds to receive the reward. Mice are held at stage 4 to ensure that the 3-second trained interval is internalized.

During each trial, one target zone is correct, but the pattern is pseudorandomized and bias-corrected to encourage mice to explore both target options and reduce perseveration. A pause in either zone (rewarded or non-rewarded) triggers the audible tone, which terminates if the mouse begins to move above the threshold speed. During training, both sides have equal (bias corrected) chance of being rewarded, and the mouse is free to check both targets during each trial. Thus, if a reward is not produced after waiting at one target for a specific interval of time, the mouse can leave to check the alternative target. Since the tone continues while the mouse remains at either target, the time that they begin moving represents an internal estimate of the trained interval.

During training, mice gradually learn to leave a target zone when they estimate the 3-second pause interval has passed without reward, indicating a timed pause in the non-rewarded zone. In practice, mice often attempt to restart the trial and return to the same target zone for another pause multiple times before checking the alternate target. Each such attempt, or a timed-pause prior to leaving a target zone, is considered a sub-trial. The average pause time on the non-rewarded side is an estimate of the animal’s internal estimate of 3 seconds. Stops less than 1 second are discarded and not analyzed. This version of the head-fixed interval timing task did not have a time-out period or cut-off, so mice could pause for extended amounts of time. However, we defined pauses greater than 9 seconds as non-engaged and are not analyzed; these types of pauses comprise ∼20% of all pauses greater than 1 second. The ratio of these durations, 3-second trained interval and 9-second cut-off, is similar to the freely-moving interval timing task (6-second trained interval and 18-second cut-off), other versions of the freely-moving interval timing task (Balci, Papachristos, et al., 2008; DeCoteau & Fox, 2022; Tosun et al., 2016), as well as peak interval timing tasks in which subjects can continue to respond up to three times the trained interval during probe trials (Meck, 2005; Rakitin et al., 1998).

### Neuronal recordings

Neuronal ensemble activity in the prefrontal cortex was recorded during interval timing sessions using a multielectrode system (Open Ephys, Atlanta, GA). Of note, separate recording sessions from the operant based freely-moving interval timing task in these mice are included in previous reports (Ding et al., 2026; Weber et al., 2025). Spike sorting was performed offline using Plexon Offline Sorter™ (Plexon, Dallas, TX), which employed principal component analysis (PCA) and waveform morphology to remove artifacts and classify single units. Two freely-moving interval timing task sessions were sorted together as in our prior work (Bova et al., 2026; Weber et al., 2025). Due to the head-fixed interval timing task training protocol and the amount of time between freely-moving and head-fixed recording sessions (average 10 months, which was constrained by availability of head-fixed behavioral equipment), we did not attempt to sort individual neurons across the different interval timing tasks. Accordingly, the neuronal ensembles from each interval timing task are considered statistically independent.

Single units were identified based on three criteria: (1) consistent waveform shape, (2) distinct clustering in PCA space, and (3) a minimum 2-ms refractory period in the inter-spike interval histogram. Spike activity was quantified using kernel density estimation of firing rates over a window spanning from 4 seconds before to 22 seconds after trial onset in the freely-moving interval timing task and 4 seconds before to 13 seconds after trial onset in the head-fixed interval timing task. Firing rates were calculated in 0.1-second bins using a Gaussian kernel with a 1-second bandwidth for principal component analyses and a 0.2-second bandwidth for general linear models.

Consistent with our previous studies (Bova et al., 2026; Bruce et al., 2021; Emmons et al., 2017; Y.-C. Kim & Narayanan, 2019; Weber et al., 2025), time-related ramping activity was defined as a monotonic change in firing rate across the entire interval. For each neuron, generalized linear models (GLMs; *fitglme* in MATLAB) were used to estimate the average firing rate slope and its standard error across trials. Time-related ramping was determined by a significant linear fit, assessed via *ANOVA* in MATLAB. Only neurons with *p* < 0.05 after Bonferroni-Holm correction for multiple comparisons were classified as exhibiting time-related ramping.

### Time-frequency analyses

Our previous studies have characterized low-frequency prefrontal cortical oscillations at ∼4 Hz during interval timing (Emmons et al., 2016; Parker, Ruggiero, et al., 2015; Weber et al., 2025). Following similar methods, we computed spectral power by convolving the fast Fourier transform (FFT) of single-trial data with the FFT of a family of complex Morlet wavelets, defined as Gaussian-windowed complex sine waves. These wavelets spanned 1–40 Hz in 100 logarithmically spaced frequency steps, with the number of cycles per wavelet increasing linearly from 3 to 12 cycles across the frequency range. The inverse FFT was then used to extract the time-frequency representation.

To facilitate comparison across frequencies and treatment conditions, spectral power was normalized to a decibel (dB) scale using the formula: 10 × log□□[power(t)/power(baseline)]. Given the self-initiated nature of the interval timing task, the pre-trial baseline was defined as the period from −3 to −2 seconds before trial onset (Weber et al., 2025). In the head-fixed interval timing task, because we are analyzing sub-trials as described above, the pre-trial baseline was defined as −0.3 to −0.2 seconds before sub-trial onset. As in previous work, we used two-tailed *t*-tests to identify significant differences between treatment conditions, with a minimum cluster size of 500 contiguous pixels used to control for multiple comparisons (Parker, Chen, et al., 2015; Singh et al., 2023; Weber et al., 2025).

### Statistics

All data were analyzed with custom written MATLAB code. As in our previous work, we focused on two primary behavioral measures from the freely-moving interval timing task: 1) the mean switch response time of all trials and 2) the switch response time coefficient of variation (CV) of all switch response times. CV was calculated as the standard deviation divided by the mean and is expressed as a percentage. We calculated parallel metrics from the head-fixed interval timing task, using all non-rewarded pause times between 1-9 seconds. Stops that lasted less than 1 second or more than 9 seconds were not analyzed as described above.

Neuronal ensembles from the different interval timing tasks are considered statistically independent. Differences in principal component analyses from each task were analyzed using one-way repeated measures ANOVA (*lmer* and *anova*). Chi-squared tests (*crosstab*) were used to compared proportions of neuronal ensembles that display time-dependent ramping activity, and differences between time-related ramping neurons across the interval timing tasks were analyzed using two-sample t-tests (*ttest2*).

## RESULTS

### Mice display time-based behavior in freely-moving and head-fixed interval timing tasks

We sought to further elucidate the neural mechanisms of interval timing by comparing two time-based tasks within individual animals: 1) a freely-moving interval timing task that requires mice to move and perform a motor response and 2) a head-fixed interval timing task that requires mice to hold still during the timed interval. In the freely-moving interval timing task, we define the switch response time as the moment the mouse exits the short nosepoke prior to responding at the long nosepoke (Fig. 1A), which is guided by internal representations of how much time has passed. The mean switch response density peaks after 6 seconds around 10.4 seconds (Fig. 1B) and the mean cumulative probability of a switch indicates that only 14.5% of switch responses occur before 6 seconds (Fig. 1C). Mean switch response times (10.39 ± 0.38 seconds) and switch time coefficient of variation (33.90 ± 1.96%) are shown in Fig 1D and 1E, respectively.

**Figure 1.**
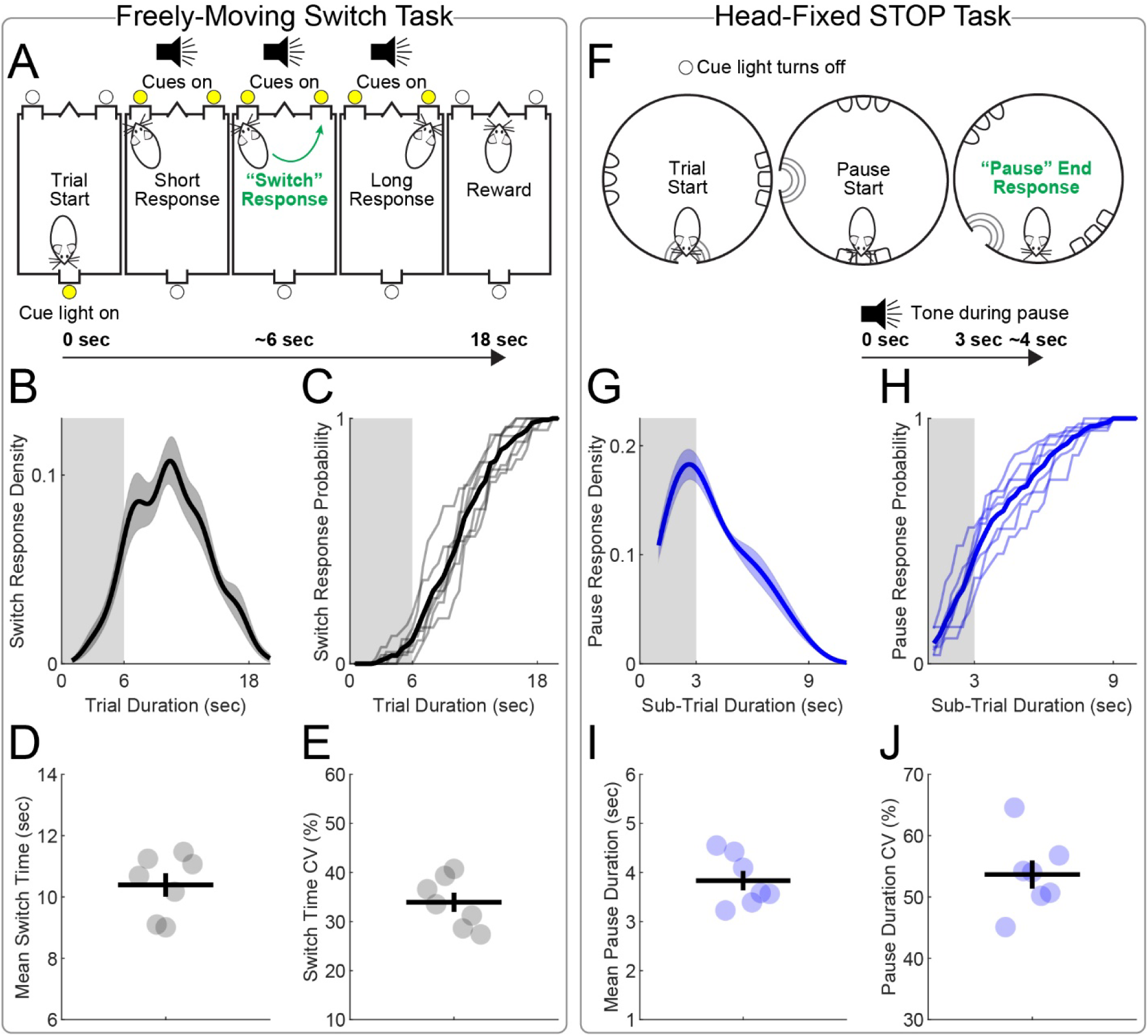
Comparison of freely-moving and head□fixed interval timing behaviors within the same animals. **A)** Freely-moving interval timing schematic showing trial initiation, entry into the short nosepoke, and the switch response defined as the moment the mouse exits the short port before moving to the long port, reflecting internally generated estimates of elapsed time. **B)** Switch response densities showed a clear peak well after the 6-second short interval. **C)** Cumulative switch response probabilities indicated that only a minority of responses occurred before the 6-second short interval. **D–E)** Mean switch times and switch time coefficient of variance (CV). **F)** Head□fixed interval timing task schematic in which the timed response is a pause: mice hold still in the absence of external cues marking the trained interval. **G)** Pause duration densities for engaged pauses showed a clear peak near the trained interval. **H)** Cumulative pause probabilities indicated that many pauses extended beyond the trained interval. **I–J)** Mean pause durations and pause CV. Data from 7 mice trained on both tasks.

In the head-fixed interval timing task, the timed response is instead a pause in which the mouse stands still while awaiting a potential reward (Fig. 1F). Importantly, because this task does not provide external cues as to when 3 seconds have passed, mice can continue to hold still beyond the 3-second trained interval when on the unrewarded target. Accordingly, response densities for engaged pauses, defined as non-rewarded pauses between 1-9 seconds, show that pause durations peak at 2.6 seconds (Fig. 1G). However, the cumulative probability of a pause response indicates that 51% of these pause responses occur after 3 seconds (Fig. 1H). Mean pause response times (3.83 ± 0.20 seconds) and pause time coefficient of variation (53.65 ± 2.31%) are shown in Fig 1I and 1J, respectively.

### Prefrontal cortex neuronal ensembles exhibit time-related ramping in both tasks

We recorded prefrontal cortex neuronal ensembles from 7 mice during the freely-moving interval timing task, using methods described in our previous work (Bova et al., 2026; Weber et al., 2025). We sorted and matched 123 neurons across two freely-moving interval timing task sessions. Neuron waveform shape across sessions was highly correlated (R = 0.997 ± 0.001; data not shown). After completing the freely-moving interval timing task, the same mice were then trained on the head-fixed interval timing task. We sorted 95 neurons from 6 mice during the head-fixed interval timing task; one mouse did not have any neurons that met criteria for inclusion; however, this subject was included for behavior and time-frequency spectral analyses.

We sought to characterize neuronal activity across the freely-moving interval timing task and head-fixed interval timing task. We used data-driven principal component analysis (PCA) to quantify patterns of neuronal activity within each task. Similar to our previous work, we generated peri-event time histograms (PETHs) using kernel density estimates and z-score normalization of firing rates between −4 and 22 seconds around trial start for the freely-moving interval timing task (Fig. 2A). For the freely-moving interval timing task, PCA identified time-dependent linear changes in z-scored firing rate over the 18-second interval as PC1, explaining ∼35% of the variance among neurons (Fig. 2B), which is similar to our prior work (Bova et al., 2026; Weber et al., 2025). PC magnitudes from the freely-moving interval timing task were significantly different (one-way ANOVA: F_(2, 366)_ = 19.0922, *p* = 1.2972 × 10^-8^; Fig. 2C). PC1 |scores| were significantly greater than both PC2 |scores| (*p* = 0.02) and PC3 |scores| (*p* = 5.53 × 10^-9^), and PC2 |scores| were significantly greater than PC3 |scores| (*p* = 0.002).

**Figure 2.**
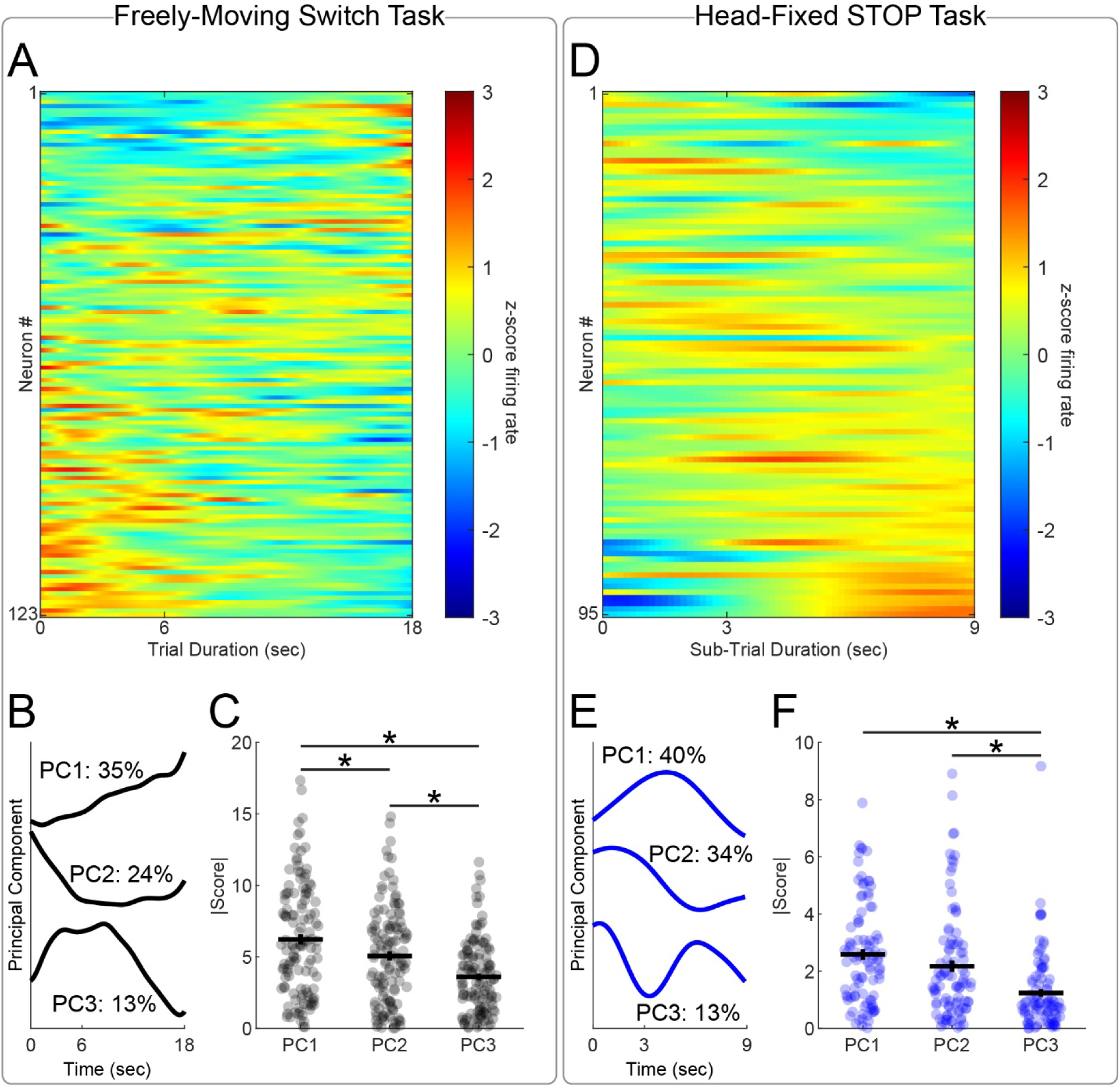
Prefrontal cortex ensembles display time-dependent activity. **A)** Peri-event time histogram (PETH) showing single neurons (n = 123) from 7 mice performing the freely-moving interval timing task. Each row represents a PETH across all switch trials, binned at 0.1 seconds, smoothed using kernel density estimates, and z scored. Colors indicate z-scored firing rates. **B)** Principal component analysis (PCA) of prefrontal cortex ensembles during the freely-moving interval timing task identified PC1 as time-dependent linear changes in z-scored firing rate over the 18-second interval and explained 35% of population variance. **C)** PC |scores| from prefrontal cortex ensembles during the freely-moving interval timing task; each dot represents |score| from a single neuron; horizontal line represents the mean, and the vertical line represents standard error. * indicates *p* < 0.05 via Bonferroni correction for multiple comparisons. **D)** PETH showing single neurons (n = 95) from 6 mice. Each row represents a PETH across all pause trials, binned at 0.1 seconds, smoothed using kernel density estimates, and z scored. Colors indicate z scored firing rates. **E)** PCA of prefrontal cortex ensembles during the head-fixed interval timing task identified PC1 as time-dependent linear changes in z-scored firing rate to approximately 3-4 seconds and explained 40% of population variance. **F)** PC |scores| from prefrontal cortex ensembles during the head-fixed interval timing task; each dot represents |score| from a single neuron; horizontal line represents the mean, and the vertical line represents standard error. * indicates *p* < 0.05 via Bonferroni correction for multiple comparisons.

For the head-fixed interval timing task, we generated PETHs between −4 and 13 seconds around pause start (Fig. 2D). PCA identified a pattern of activity in which time-dependent linear changes in z-scored firing rate is evident until approximately 3-4 seconds and explained ∼40% of the variance (Fig. 2E). Similar to the freely-moving interval timing task, PC magnitudes from the head-fixed interval timing task were significantly different (one-way ANOVA: F_(2, 282)_ = 15.73, *p* = 3.36 × 10^-7^; Fig. 2F). PC1 |scores| were not significantly different than PC2 |scores| (*p* = 0.21). However, both PC1 |scores| and PC2 |scores| were significantly greater than PC3 |scores| (*p* = 1.3 × 10^-7^ and *p* = 0.0004, respectively).

We then turned to general linear models (GLMs) that quantify neuronal ramping activity on a trial-by-trial basis to further interrogate ramping dynamics across interval timing tasks. Since PCA suggested a pattern of ramping activity until 3-4 seconds in the head-fixed interval timing task, we focused on quantifying patterns of ramping activity until the end of the trained interval in each task (i.e., 6 seconds in the freely-moving interval timing task and 3 seconds in the head-fixed interval timing task). Example neurons that display time-dependent ramping activity over the trained interval are shown in Fig. 3A and Fig. 3B for the freely-moving interval timing task and head-fixed interval timing task, respectively. For each neuron during the freely-moving interval timing task, we constructed a model in which *Firing Rate* is the response variable and *Time* in the interval is the predictor variable. *Responses* (nosepoke responses during the trial) is included as a regressor for movement. Identical models were generated for each neuron during the head-fixed interval timing task; however, *Responses* was omitted as a regressor since this task explicitly requires a lack of movement. For each neuron in each task, we obtained the corrected *p* value for main effect of time, the trial-by-trial slope estimate, and the estimated trial-by-trial standard error. This analysis includes trial-by-trial neuronal firing rates and is not influenced by trial averaging. Ramping neurons were defined by GLM fits and a Bonferroni-Holm corrected significant linear change in firing rate over the trained interval. This analysis identified that 40% (49/123) of prefrontal cortex neurons display time-dependent ramping activity over the first 6 seconds of the freely-moving interval timing task. Separate GLMs identified that 18% (17/95) of neurons display ramping activity over the 3-second interval during the head-fixed interval timing task, which is significantly less frequent than the freely-moving interval timing task (Χ^2^ = 12.23; *p* = 0.0005; Fig. 3C).

**Figure 3.**
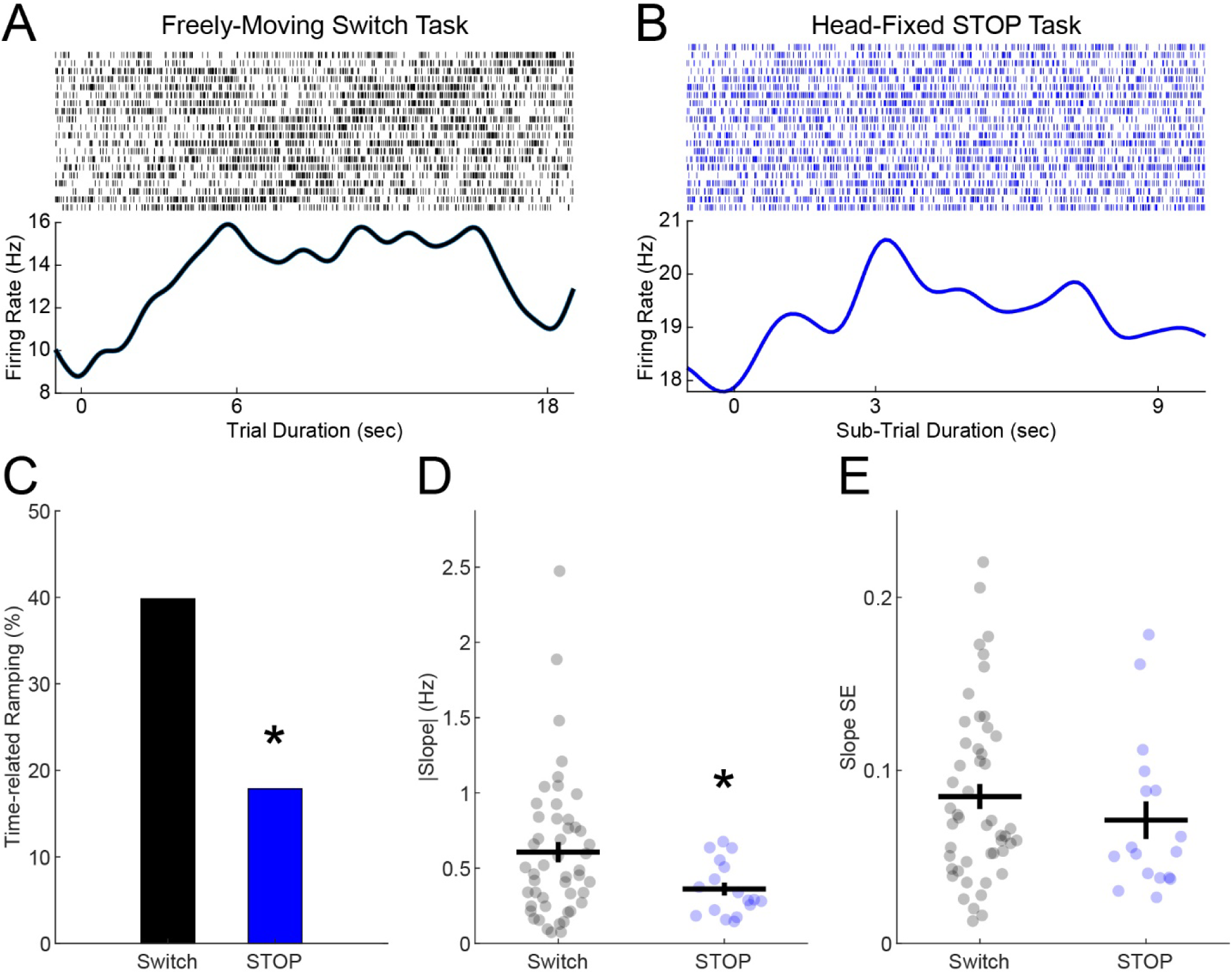
Prefrontal cortex neurons display time-dependent ramping activity over the trained interval. Peri-event rasters (top) and estimated mean firing rate (bottom) of neurons that display time-related ramping activity during the **(A)** 6-second trained interval in the freely-moving interval timing task and **(B)** 3-second trained interval in the head-fixed interval timing task. **C)** Ramping activity was more common in the freely-moving interval timing task compared to the head-fixed interval timing task. **D)** Ramping neurons displayed a greater linear |slope| in the freely-moving interval timing task compared to ramping neurons in the head-fixed interval timing task, but **(E)** these neurons did not display differences in slope standard error (SE).

While there is a difference in the relative percentage of neurons that display time-related ramping activity, we sought to also determine whether there are differences in the rate of change (i.e., slope) or how variable a neuron ramps over multiple trials. We tested these ideas by comparing the GLM estimate of trial-by-trial slope and trial-by-trial standard error from time-related ramping neurons during the freely-moving interval timing task and head-fixed interval timing task. We found that trial-by-trial slope was significantly different (freely-moving: 0.6069 ± 0.0671 Hz vs. head-fixed; 0.3617 ± 0.0435 Hz; t_(1,64)_ = 2.0920; *p* = 0.0404; Fig. 3D) but trial-by-trial standard error was not (freely-moving: 0.0849 ± 0.0072 vs. head-fixed; 0.0712 ± 0.0109; t_(1,64)_ = 0. 9927; *p* = 0.3246; Fig. 3E).

Across both tasks, each mouse indicates its internal representation of how much time has passed at one target — the switch response in the freely-moving interval timing task and the end of the pause in the head-fixed interval timing task. We sought to determine if ramping also occurs in the time leading up to the timed decision. Specifically, we focused on 3 seconds prior to the timed decision in each task. Example neurons that display time-related ramping activity over the 3 seconds before the timed decision are shown in Fig. 4A and Fig. 4B for the freely-moving interval timing task and head-fixed interval timing task, respectively. Interestingly, using similar GLMs as above, we found that 33% of neurons ramp over the 3 seconds before the timed decision in both the freely-moving interval timing task (41/123 neurons) and the head-fixed interval timing task (31/95 neurons); these percentages are not significantly different (Χ^2^ = 0.0119; *p* = 0.9130; Fig. 4C). We also found that there was no difference in either trial-by-trial slope (freely-moving interval timing: 0.3193 ± 0.0480 Hz vs. head-fixed interval timing task; 0.3775 ± 0.0691 Hz; t_(1,70)_ = 0.7130; *p* = 0.4782; Fig. 4D) or trial-by-trial standard error (freely-moving interval timing task: 0.0806 ± 0.0081 vs. head-fixed interval timing task; 0.0662 ± 0.0087; t_(1,70)_ = 1.2008; *p* = 0.2339; Fig. 4E). Taken together, these data show that ramping activity is present during interval timing, even when mice are actively holding still and not moving, providing new insight into prefrontal neuronal ramping activity.

**Figure 4.**
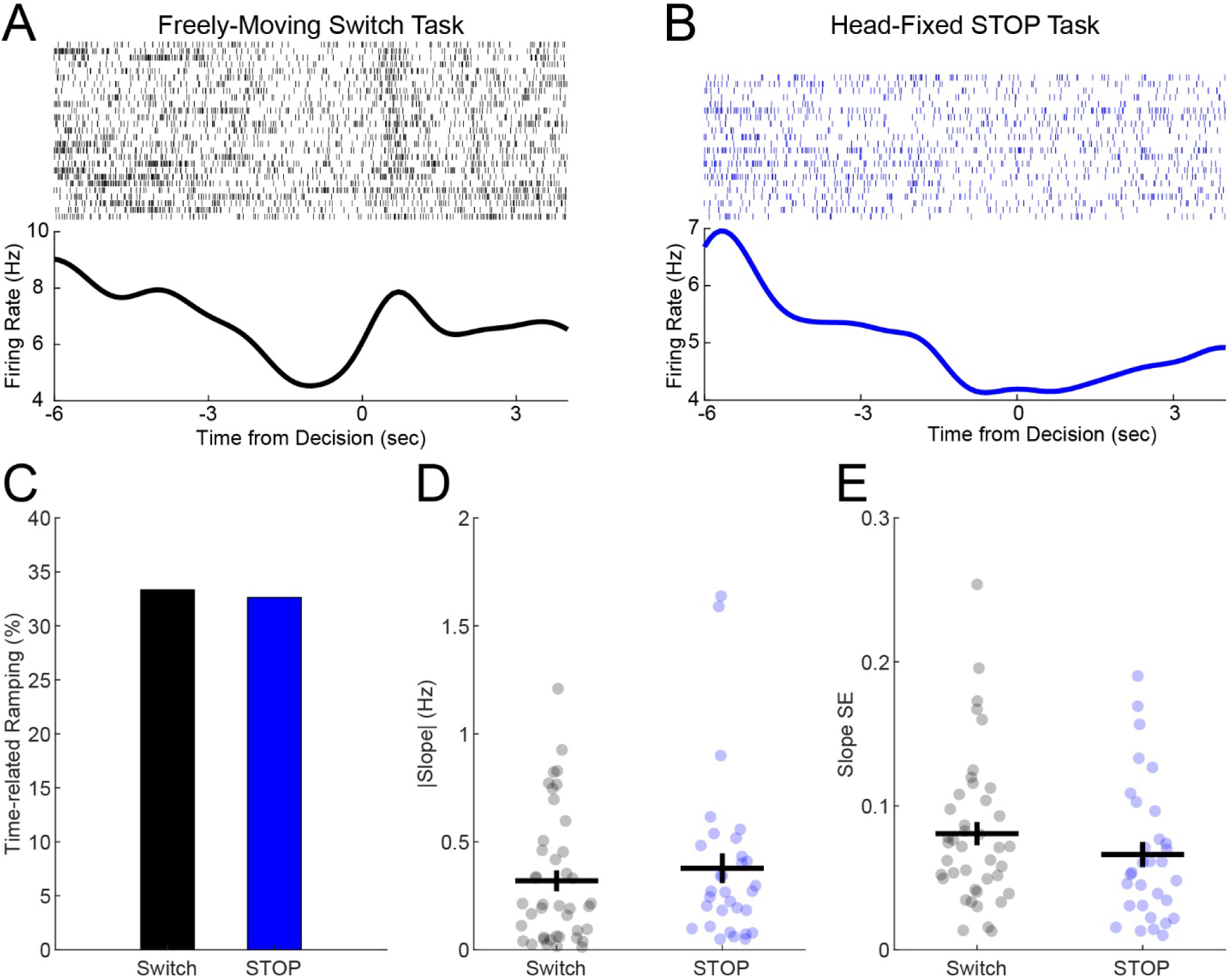
Prefrontal cortex neurons display time-dependent ramping activity prior to the timed decision. Peri-event rasters (top) and estimated mean firing rate (bottom) of neurons that display time-related ramping activity 3 seconds prior to the timed decision in **(A)** the freely-moving interval timing task and **(B)** the head-fixed interval timing task. **C)** The proportion of neurons that display ramping activity over the 3 seconds before the decision did not differ between the two tasks. **D)** Linear |slope| and **(E)** standard error (SE) did not differ between ramping neurons in the freely-moving interval timing task compared to ramping neurons in the head-fixed interval timing task.

### Interval timing triggers low-frequency prefrontal cortical rhythms

An important feature of interval timing is that this behavior elicits low-frequency prefrontal “theta” rhythms that recruit neural mechanisms crucial for cognitive control (Cavanagh & Frank, 2014). These low-frequency bursts of activity are dampened in human patients and rodent models with interval timing deficits (Y.-C. Kim et al., 2017; Parker, Chen, et al., 2015; Singh et al., 2021, 2023; Weber et al., 2025). Consistent with our prior work, we found a burst of low-frequency rhythms between 2-4 Hz and 4-7 Hz at the start of trials during the freely-moving interval timing task (Fig. 5A). We also found similar patterns of low-frequency activity at the start of trials during the head-fixed interval timing task (Fig. 5B).

**Figure 5.**
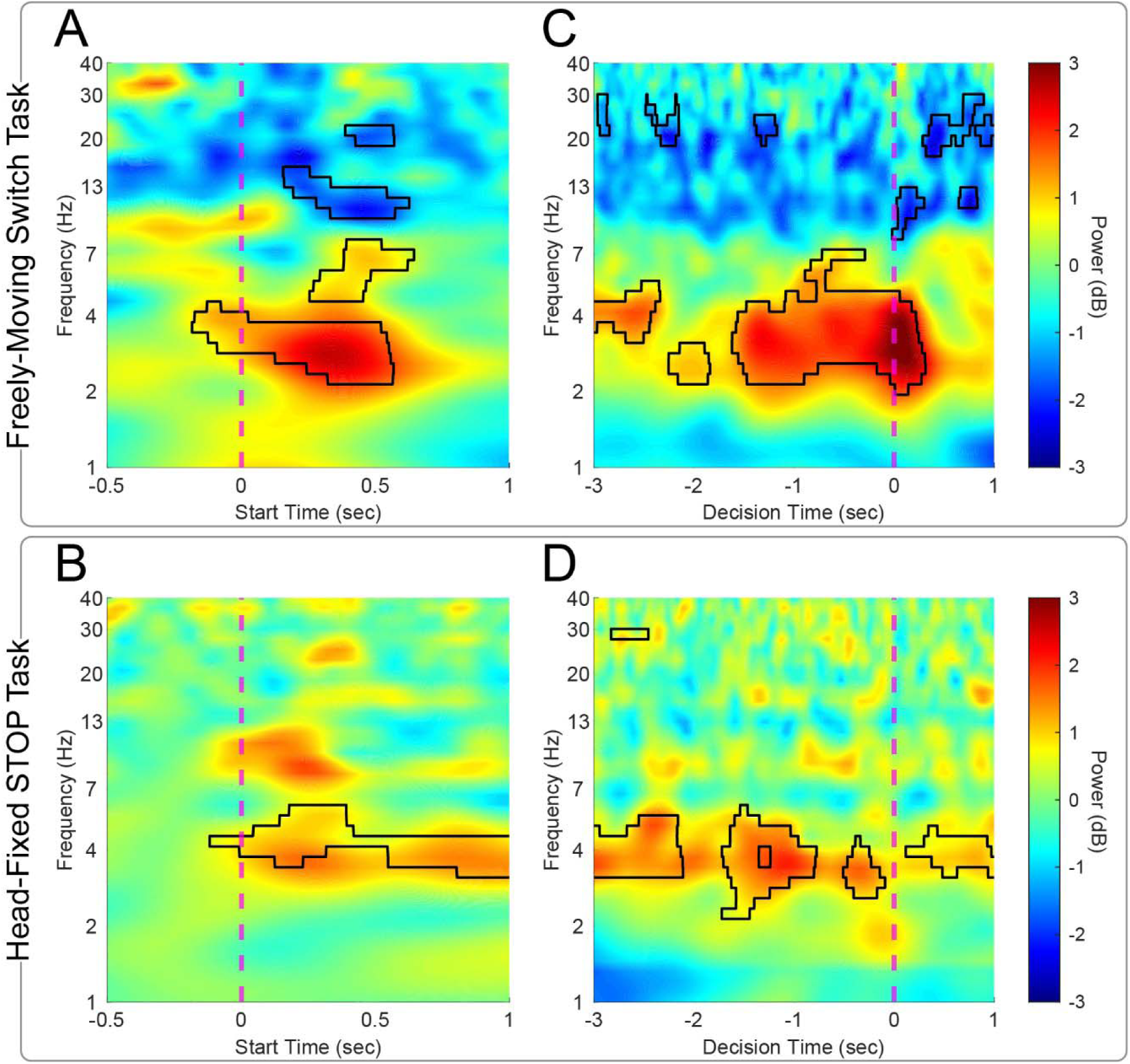
Low-frequency cortical rhythms during interval timing. Time-frequency analysis showed a burst of low-frequency ∼4 Hz power at trial start during both the freely-moving interval timing task **(A)** and head-fixed interval timing task **(B)**. Similar bursts of activity are observed around the timed decision in both tasks **(C-D).** Areas outlined with solid black lines indicate clusters sizes of at least 500 pixels with *p* < 0.05 via t-test, representing a significant difference from baseline.

Further, we were also interested in low-frequency rhythms in anticipation of the timed decision, corresponding to the switch in freely-moving interval timing and the end of the pause in head-fixed interval timing. We again found a prominent burst of low-frequency activity, which is sustained for more than 1 second before the timed decision in both tasks (Fig. 5C-D). These results further support the hypothesis that 4 Hz activity is engaged by interval timing at trial start and in anticipation of a timed decision and is engaged regardless of the specific motor program.

## DISCUSSION

Rodent interval timing tasks typically require a motor response, so we tested the hypothesis that prefrontal cortex neuronal ramping activity and evoked 4 Hz activity would occur in interval timing tasks independent of the timed motor response. We trained mice to perform a freely-moving version of interval timing where subjects reported elapsed time by switching nosepokes and a head-fixed version of interval timing where animals had to hold their entire body still for a pause of 3 seconds. Strikingly, we found prominent prefrontal ramping activity and evoked 4 Hz activity in both tasks of generally equivalent magnitude. Taken together, these data suggest that ramping activity and 4 Hz activity are not linked to specific motor programs and are more likely related to actively estimating a temporal interval.

Neuronal ramping activity is observed ubiquitously in the prefrontal cortex during tasks with a temporal interval (Merchant & Averbeck, 2017; Narayanan, 2016; Wang et al., 2018) and consistent with drift-diffusion models of accumulating evidence (Simen et al., 2011). However, one hypothesis is that neuronal ramping activity may also reflect motor sequencing. To address this, one recent study by our laboratory measured movement during interval timing as well as reward anticipation during a Pavlovian conditioning task, and found that neither movement nor reward anticipation could fully account for ramping activity (Bova et al., 2026). However, most interval timing tasks require movement, and it is unknown whether movement is required for time-related neuronal activity in the prefrontal cortex. To test this hypothesis, we developed a novel head-fixed interval timing task in which animals must estimate time while actively inhibiting movements, in contrast to freely-moving interval timing, where mice must make movements to estimate time. While different animals and species exhibit prefrontal ramping under a wide range of conditions, including decision-making tasks, hold-tasks, reaction-time tasks, and even working-memory tasks (Narayanan et al., 2013; Niki & Watanabe, 1976; Sheth et al., 2012; Tanji & Evarts, 1976), to our knowledge, few studies have recorded from the same prefrontal networks under two conditions and directly compared ramping signals.

We also report that ∼4 Hz theta activity is observed in both tasks. This activity has been attributed to cognitive control when effort is required such as at adjudicating conflict, punishment, or adjusting after an error (Cavanagh & Frank, 2014). However, one of the most reliable epochs of this signal in prefrontal networks is at the start of the trial, which requires engagement of task-specific activity. Our past work has shown that this 4 Hz activity can be influenced by past trials and required dopamine, and is altered in human diseases such as Parkinson’s disease and schizophrenia (Laubach et al., 2015; Parker et al., 2017; Singh et al., 2023).

We find that 4 Hz activity is present at the beginning of the trial in both freely-moving and head-fixed interval timing, although it is sustained in head-fixed interval timing, potentially because animals are required to stay engaged to hold still. We also present evidence consistent with prior work in rats that ∼4 Hz activity is apparent prior to when the decision to respond is made. This is remarkable because in the freely-moving interval timing task, the mouse is required to move while timing the interval, compared to the head-fixed interval timing task in which the mouse is, by definition, not moving during the timed interval.

While our work clearly shows prefrontal neuronal ramping and ∼4 Hz activity in two distinct interval timing contexts, there are several limitations. First, because our mice required extensive retraining, we could not hold neurons between tasks and make explicit comparisons across the same individual neurons. We also cannot rule out that some differences in neuronal activity observed between the tasks are secondary to aging. We therefore focus on the intriguing similarities found between the tasks, and future studies of the head-fixed task in younger mice will be useful for evaluating differences. Second, the molecular and circuit identities of ramping neurons are not fully known, and remain an area of active research (Ding et al., 2026); however, head-fixed interval timing tasks may help identify these neurons individually and help determine what specific interval these neurons are encoding. Third, while we have extensive data on how freely-moving interval timing tasks respond to manipulations relevant to neurodegenerative disease and drugs (Stutt et al., 2024; Weber et al., 2023, 2025), these perturbations have yet to be established for head-fixed interval timing.

In conclusion, we found that ramping activity and ∼4 Hz theta activity is observed in the prefrontal cortex of the same mice during two tasks that require opposite and distinct motor activity, providing evidence that these features of prefrontal activity are related to timing intervals rather than task-specific movements or strategies. Future work will further establish how these aspects of prefrontal activity work together to generate flexible timed behavior and may malfunction in human brain disease.

## ACKNOWLEDGMENTS

Authorship

MAW, JR, PJB, YK, NSN, and GMA designed the experiments. MAW, JR, and KG performed all experiments. MAW, JR, KG, ASB, YK, NSN, and GMA performed all statistical analyses. MAW, JR, NSN, and GMA wrote the manuscript, and all authors reviewed and revised the manuscript.

## REFERENCES

Albert, M. S. (1996). Cognitive and neurobiologic markers of early Alzheimer disease. Proceedings of the National Academy of Sciences of the United States of America, 93(24), 13547–13551. 10.1073/pnas.93.24.13547

Andreasen, N. C. (1999). Understanding the causes of schizophrenia. The New England Journal of Medicine, 340(8), 645–647. 10.1056/NEJM199902253400811

Baddeley, A. D., Baddeley, H. A., Bucks, R. S., & Wilcock, G. K. (2001). Attentional control in Alzheimer’s disease. Brain: A Journal of Neurology, 124(Pt 8), 1492–1508. 10.1093/brain/124.8.1492

Balci, F., Ludvig, E. A., Gibson, J. M., Allen, B. D., Frank, K. M., Kapustinski, B. J., Fedolak, T. E., & Brunner, D. (2008). Pharmacological manipulations of interval timing using the peak procedure in male C3H mice. Psychopharmacology, 201(1), 67–80. 10.1007/s00213-008-1248-y

Balci, F., Papachristos, E. B., Gallistel, C. R., Brunner, D., Gibson, J., & Shumyatsky, G. P. (2008). Interval timing in genetically modified mice: A simple paradigm. Genes, Brain and Behavior, 7(3), 373–384. 10.1111/j.1601-183X.2007.00348.x

Bova, A., Weber, M. A., Spicer, M. M., Khandelwal, V., Volkman, R. A., McMurrin, M., Kryca, J., Sivakumar, K., Kirkpatrick, B. Q., Zhang, Q., & Narayanan, N. S. (2026). Timing, movement, and reward contributions to prefrontal and striatal ramping activity. The Journal of Neuroscience: The Official Journal of the Society for Neuroscience, e1493252026. 10.1523/JNEUROSCI.1493-25.2026

Bruce, R. A., Weber, M. A., Volkman, R. A., Oya, M., Emmons, E. B., Kim, Y., & Narayanan, N. S. (2021). Experience-related enhancements in striatal temporal encoding. The European Journal of Neuroscience, 54(3), 5063–5074. 10.1111/ejn.15344

Bruce, R. A., Weber, M., Bova, A., Volkman, R., Jacobs, C., Sivakumar, K., Stutt, H., Kim, Y., Curtu, R., & Narayanan, K. (2025). Complementary cognitive roles for D2-MSNs and D1-MSNs during interval timing. eLife, 13, RP96287. 10.7554/eLife.96287

Buhusi, C. V., & Meck, W. H. (2005). What makes us tick? Functional and neural mechanisms of interval timing. Nature Reviews Neuroscience, 6(10), 755–765. 10.1038/nrn1764

Caselli, L., Iaboli, L., & Nichelli, P. (2009). Time estimation in mild Alzheimer’s disease patients. Behavioral and Brain Functions, 5(1), 32. 10.1186/1744-9081-5-32

Cavanagh, J. F., & Frank, M. J. (2014). Frontal theta as a mechanism for cognitive control. Trends in Cognitive Sciences, 18(8), 414–421. 10.1016/j.tics.2014.04.012

DeCoteau, W. E., & Fox, A. E. (2022). Timing and Intertemporal Choice Behavior in the Valproic Acid Rat Model of Autism Spectrum Disorder. Journal of Autism and Developmental Disorders, 52(6), 2414–2429. 10.1007/s10803-021-05129-y

Diamond, A. (2013). Executive Functions. Annual Review of Psychology, 64(1), 135–168. 10.1146/annurev-psych-113011-143750

Ding, X., Weber, M. A., Butler, T. C., Bova, A. S., Guerrero, S. G., Hunter, C. M., Cole, R. C., Stutt, H. R., McMurrin, M. S., Spicer, M. M., Conlon, M. M., Heiney, S. A., Kim, Y., Resch, J. M., & Narayanan, N. S. (2026). Distinct activity in prefrontal projections promotes temporal control of action. Proceedings of the National Academy of Sciences, 123(19), e2538059123. 10.1073/pnas.2538059123

Dolan, S. L., Bechara, A., & Nathan, P. E. (2008). Executive dysfunction as a risk marker for substance abuse: The role of impulsive personality traits. Behavioral Sciences & the Law, 26(6), 799–822. 10.1002/bsl.845

Emmons, E. B., De Corte, B. J., Kim, Y., Parker, K. L., Matell, M. S., & Narayanan, N. S. (2017). Rodent Medial Frontal Control of Temporal Processing in the Dorsomedial Striatum. The Journal of Neuroscience, 37(36), 8718–8733. 10.1523/JNEUROSCI.1376-17.2017

Emmons, E. B., Ruggiero, R. N., Kelley, R. M., Parker, K. L., & Narayanan, N. S. (2016). Corticostriatal Field Potentials Are Modulated at Delta and Theta Frequencies during Interval-Timing Task in Rodents. Frontiers in Psychology, 7. 10.3389/fpsyg.2016.00459

Fuster, J. M. (2001). The prefrontal cortex—An update: Time is of the essence. Neuron, 30, 319–333. (11394996).

Gür, E., Duyan, Y. A., Arkan, S., Karson, A., & Balcı, F. (2020). Interval timing deficits and their neurobiological correlates in aging mice. Neurobiology of Aging, 90, 33–42. 10.1016/j.neurobiolaging.2020.02.021

Gür, E., Fertan, E., Kosel, F., Wong, A. A., Balcı, F., & Brown, R. E. (2019). Sex differences in the timing behavior performance of 3xTg-AD and wild-type mice in the peak interval procedure. Behavioural Brain Research, 360, 235–243. 10.1016/j.bbr.2018.11.047

Henke, J., Bunk, D., Von Werder, D., Häusler, S., Flanagin, V. L., & Thurley, K. (2021). Distributed coding of duration in rodent prefrontal cortex during time reproduction. eLife, 10, e71612. 10.7554/eLife.71612

Hinton, S. C., & Meck, W. H. (2004). Frontal–striatal circuitry activated by human peak-interval timing in the supra-seconds range. Cognitive Brain Research, 21(2), 171–182. 10.1016/j.cogbrainres.2004.08.005

Keyes, A. L., Kim, Y., Bosch, P. J., Usachev, Y. M., & Aldridge, G. M. (2021). Stay or go? Neuronal activity in medial frontal cortex during a voluntary tactile preference task in head-fixed mice. Cell Calcium, 96, 102388. 10.1016/j.ceca.2021.102388

Kim, J., Ghim, J.-W., Lee, J. H., & Jung, M. W. (2013). Neural Correlates of Interval Timing in Rodent Prefrontal Cortex. The Journal of Neuroscience, 33(34), 13834–13847. 10.1523/JNEUROSCI.1443-13.2013

Kim, Y.-C., Han, S.-W., Alberico, S. L., Ruggiero, R. N., De Corte, B., Chen, K.-H., & Narayanan, N. S. (2017). Optogenetic Stimulation of Frontal D1 Neurons Compensates for Impaired Temporal Control of Action in Dopamine-Depleted Mice. Current Biology, 27(1), 39–47. 10.1016/j.cub.2016.11.029

Kim, Y.-C., & Narayanan, N. S. (2019). Prefrontal D1 Dopamine-Receptor Neurons and Delta Resonance in Interval Timing. Cerebral Cortex, 29(5), 2051–2060. 10.1093/cercor/bhy083

Larson, T., Khandelwal, V., Weber, M. A., Leidinger, M. R., Meyerholz, D. K., Narayanan, N. S., & Zhang, Q. (2022). Mice expressing P301S mutant human tau have deficits in interval timing. Behavioural Brain Research, 432, 113967. 10.1016/j.bbr.2022.113967

Laubach, M., Caetano, M. S., & Narayanan, N. S. (2015). Mistakes were made: Neural mechanisms for the adaptive control of action initiation by the medial prefrontal cortex. Journal of Physiology-Paris. 10.1016/j.jphysparis.2014.12.001

Malapani, C., Rakitin, B., Levy, R., Meck, W. H., Deweer, B., Dubois, B., & Gibbon, J. (1998). Coupled temporal memories in Parkinson’s disease: A dopamine-related dysfunction. Journal of Cognitive Neuroscience, 10(3), 316–331.

Meck, W. H. (2005). Neuropsychology of timing and time perception. Brain and Cognition, 58(1), 1–8. 10.1016/j.bandc.2004.09.004

Merchant, H., & Averbeck, B. B. (2017). The Computational and Neural Basis of Rhythmic Timing in Medial Premotor Cortex. The Journal of NeurosciencelJ: The Official Journal of the Society for Neuroscience, 37(17), 4552–4564. 10.1523/JNEUROSCI.0367-17.2017

Merchant, H., & De Lafuente, V. (2014). Introduction to the Neurobiology of Interval Timing. In H. Merchant & V. De Lafuente (Eds.), Neurobiology of Interval Timing (Vol. 829, pp. 1–13). Springer New York. 10.1007/978-1-4939-1782-2_1

Narayanan, N. S. (2016). Ramping activity is a cortical mechanism of temporal control of action. Current Opinion in Behavioral Sciences, 8, 226–230. 10.1016/j.cobeha.2016.02.017

Narayanan, N. S., Cavanagh, J. F., Frank, M. J., & Laubach, M. (2013). Common medial frontal mechanisms of adaptive control in humans and rodents. Nature Neuroscience, 16(12), 1888–1897. 10.1038/nn.3549

Niki, H., & Watanabe, M. (1976). Cingulate unit activity and delayed response. Brain Res, 110, 381–386. (820408).

Parker, K. L., Chen, K.-H., Kingyon, J. R., Cavanagh, J. F., & Narayanan, N. S. (2014). D1-Dependent 4 Hz Oscillations and Ramping Activity in Rodent Medial Frontal Cortex during Interval Timing. Journal of Neuroscience, 34(50), 16774–16783. 10.1523/JNEUROSCI.2772-14.2014

Parker, K. L., Chen, K.-H., Kingyon, J. R., Cavanagh, J. F., & Narayanan, N. S. (2015). Medial frontal ∼4-Hz activity in humans and rodents is attenuated in PD patients and in rodents with cortical dopamine depletion. Journal of Neurophysiology, 114(2), 1310–1320. 10.1152/jn.00412.2015

Parker, K. L., Kim, Y. C., Kelley, R. M., Nessler, A. J., Chen, K.-H., Muller-Ewald, V. A., Andreasen, N. C., & Narayanan, N. S. (2017). Delta-frequency stimulation of cerebellar projections can compensate for schizophrenia-related medial frontal dysfunction. Molecular Psychiatry, 22(5), 647–655. 10.1038/mp.2017.50

Parker, K. L., Lamichhane, D., Caetano, M. S., & Narayanan, N. S. (2013). Executive dysfunction in Parkinson’s disease and timing deficits. Frontiers in Integrative Neuroscience, 7, 75. 10.3389/fnint.2013.00075

Parker, K. L., Ruggiero, R. N., & Narayanan, N. S. (2015). Infusion of D1 Dopamine Receptor Agonist into Medial Frontal Cortex Disrupts Neural Correlates of Interval Timing. Frontiers in Behavioral Neuroscience, 9, 294. 10.3389/fnbeh.2015.00294

Penney, T. B., Meck, W. H., Roberts, S. A., Gibbon, J., & Erlenmeyer-Kimling, L. (2005). Interval-timing deficits in individuals at high risk for schizophrenia. Brain and Cognition, 58(1), 109–118.

Rakitin, B. C., Gibbon, J., Penney, T. B., Malapani, C., Hinton, S. C., & Meck, W. H. (1998). Scalar expectancy theory and peak-interval timing in humans. Journal of Experimental Psychology. Animal Behavior Processes, 24(1), 15–33.

Sheth, S. A., Mian, M. K., Patel, S. R., Asaad, W. F., Williams, Z. M., Dougherty, D. D., Bush, G., & Eskandar, E. N. (2012). Human dorsal anterior cingulate cortex neurons mediate ongoing behavioural adaptation. Nature, 488(7410), 218–221. 10.1038/nature11239

Simen, P., Balci, F., de Souza, L., Cohen, J. D., & Holmes, P. (2011). A model of interval timing by neural integration. The Journal of Neuroscience: The Official Journal of the Society for Neuroscience, 31(25), 9238–9253. 10.1523/JNEUROSCI.3121-10.2011

Singh, A., Cole, R. C., Espinoza, A. I., Evans, A., Cao, S., Cavanagh, J. F., & Narayanan, N. S. (2021). Timing variability and midfrontal ∼4 Hz rhythms correlate with cognition in Parkinson’s disease. Npj Parkinson’s Disease, 7(1), 14. 10.1038/s41531-021-00158-x

Singh, A., Cole, R. C., Espinoza, A. I., Wessel, J. R., Cavanagh, J. F., & Narayanan, N. S. (2023). Evoked mid-frontal activity predicts cognitive dysfunction in Parkinson’s disease. Journal of Neurology, Neurosurgery & Psychiatry, 94(11), 945–953. 10.1136/jnnp-2022-330154

Stopford, C. L., Thompson, J. C., Neary, D., Richardson, A. M. T., & Snowden, J. S. (2012). Working memory, attention, and executive function in Alzheimer’s disease and frontotemporal dementia. Cortex, 48(4), 429–446. 10.1016/j.cortex.2010.12.002

Stutt, H. R., Weber, M. A., Cole, R. C., Bova, A. S., Ding, X., McMurrin, M. S., & Narayanan, N. S. (2024). Sex similarities and dopaminergic differences in interval timing. Behavioral Neuroscience, 138(2), 85–93. 10.1037/bne0000577

Tanji, J., & Evarts, E. V. (1976). Anticipatory activity of motor cortex neurons in relation to direction of an intended movement. J Neurophysiol, 39, 1062–1068. (824409).

Tosun, T., Gür, E., & Balcı, F. (2016). Mice plan decision strategies based on previously learned time intervals, locations, and probabilities. Proceedings of the National Academy of Sciences, 113(3), 787–792. 10.1073/pnas.1518316113

Wang, J., Narain, D., Hosseini, E. A., & Jazayeri, M. (2018). Flexible timing by temporal scaling of cortical responses. Nature Neuroscience, 21(1), 102. 10.1038/s41593-017-0028-6

Ward, R. D., Simpson, E. H., Kandel, E. R., & Balsam, P. D. (2011). Modeling motivational deficits in mouse models of schizophrenia: Behavior analysis as a guide for neuroscience. Behavioural Processes, 87(1), 149–156. 10.1016/j.beproc.2011.02.004

Weber, M. A., Sivakumar, K., Kirkpatrick, B. Q., Stutt, H. R., Tabakovic, E. E., Bova, A. S., Kim, Y.-C., & Narayanan, N. S. (2025). Amphetamine increases timing variability by degrading prefrontal temporal encoding. Neuropharmacology, 275, 110486. 10.1016/j.neuropharm.2025.110486

Weber, M. A., Sivakumar, K., Tabakovic, E. E., Oya, M., Aldridge, G. M., Zhang, Q., Simmering, J. E., & Narayanan, N. S. (2023). Glycolysis-enhancing α1-adrenergic antagonists modify cognitive symptoms related to Parkinson’s disease. Npj Parkinson’s Disease, 9(1), Article 1. 10.1038/s41531-023-00477-1

Wittmann, M., Leland, D. S., Churan, J., & Paulus, M. P. (2007). Impaired time perception and motor timing in stimulant-dependent subjects. Drug and Alcohol Dependence, 90(2–3), 183–192. 10.1016/j.drugalcdep.2007.03.005

